# Gene therapy targeting of AKAP6β-CaMKII signalosomes improves myocardial inflammation and heart failure in a swine model of cardiometabolic syndrome

**DOI:** 10.64898/2026.07.23.740435

**Authors:** Darla L. Tharp, Sofia M. Possidento, Jinliang Li, Abraham L. Bayer, Amira R. Amin, Pamela K. Thorne, Eryn P. Wagoner, Federico Cividini, Moriah G. Turcotte, Xueyi Li, Ying Zhu, Ramesh V. Nair, Christopher I. Murray, Vi B. Nguyen, Jennifer E. Van Eyk, Pilar Alcaide, Kimberly L. Dodge-Kafka, Craig A. Emter, Michael S. Kapiloff

## Abstract

**Background:** Cardiometabolic heart failure with preserved ejection fraction (HFpEF) is associated with systemic and cardiac inflammation and diastolic dysfunction. A-kinase anchoring protein 6β (AKAP6β) is a scaffold protein located at the cardiomyocyte outer nuclear membrane that promotes pathological cardiac remodeling via the recruitment of multiple regulatory proteins including protein kinases. In mice, adeno-associated virus (AAV) mediated expression of a peptide based upon a kinase binding domain (KBD) within AKAP6β inhibited the development of heart failure due to chronic pressure overload. Whether KBD expression can also inhibit the development of cardiometabolic heart failure is unknown, and if so, the mechanism of KBD action in HFpEF has yet to be explored.

**Methods:** The efficacy of a cardiotropic self-complementary AAV gene therapy that expresses the AKAP6β KBD peptide (AAV9sc.KBD) was tested in a female Ossabaw swine model of cardiometabolic syndrome and HFpEF. Single nucleus and bulk RNA sequencing of swine heart tissue and immunoprecipitation-mass spectrometry, live cell imaging, and biochemical assays using primary rat cardiomyocytes were employed to study KBD mechanism of action.

**Results:** AAV9sc.KBD inhibited the development of diastolic dysfunction and heart failure in the Ossabaw model, without negatively impacting systolic function. The improvement in cardiac phenotype was associated with decreased T-cell myocardial infiltrates and partial reversal of pathological gene expression. An unbiased interactome study revealed that the KBD peptide binds Ca^2+^/calmodulin-dependent protein kinase II (CaMKII), identifying CaMKII as a new AKAP6β binding partner. Perinuclear CaMKII activity detected by live cell imaging required AKAP6β expression and was inhibited by KBD expression. In addition, the CaMKII substrate Inhibitor of NF-κB Kinase β (IKKβ) bound AKAP6β. IKK phosphorylation in the Ossabaw model and in myocytes was inhibited by KBD expression, and NF-κB nuclear translocation in myocytes was dependent upon AKAP6β-CaMKII protein complex formation. AAV9sc.KBD treatment inhibited cardiomyocyte NF-κB-dependent gene expression in the Ossabaw model.

**Conclusions:** Regulated by perinuclear AKAP6β-CaMKII signalosomes, NF-κB pro-inflammatory gene expression in cardiomyocytes participates in a positive feedback loop with cardiac inflammation promoting HFpEF. Proof-of-concept is provided in a large animal model that gene therapy-based cardiomyocyte expression of the KBD peptide will prevent cardiac dysfunction in cardiometabolic syndrome.

**Clinical Perspective:** *What is new:*

- The cardiomyocyte-selective gene therapy AAV9sc.KBD, which targets signalosomes organized by the scaffold protein AKAP6β, is shown to inhibit myocardial T-cell infiltration and improve cardiac structure and function in a large animal model of cardiometabolic HFpEF.
- The AKAP6β KBD peptide is shown to bind and inhibit the function of CaMKII.
- CaMKII and IKKβ are shown to participate in perinuclear AKAP6β signalosomes, where they regulate activation of the NF-κB pro-inflammatory gene regulatory pathway.

*Clinical implications:*

- Proof-of-concept for a novel strategy for the treatment of HFpEF is provided, intracellular expression by a cardiomyocyte-selective gene therapy vector of an inhibitory peptide, which will inhibit compartmentalized intracellular signal transduction.
- In conjunction with previous studies in small rodents, the new data obtained in Ossabaw swine support clinical translation of the AAV9sc.KBD gene therapy.

## Introduction

Heart failure with preserved ejection fraction (HFpEF) is a syndrome representing over half of heart failure patients, with increased prevalence in post-menopausal women.^1^ A spectrum of systemic co-morbidities drive HFpEF, resulting in a heterogeneity of phenotypes. Characterized by decreased cardiac reserve under stress, diastolic dysfunction, left atrial enlargement, obesity, hypertension, diabetes, and dyslipidemia, cardiometabolic HFpEF is the most common HFpEF endotype and incurs particularly high morbidity.^2^ Markers of systemic inflammation are especially elevated in cardiometabolic HFpEF,^2^ resulting in myocardial inflammation that has been described as “outside-in.”^3^ For example, in a mouse model of cardiometabolic HFpEF, pro-inflammatory metabolic co-morbidities induced systemic T-cell dysfunction and expansion, followed by cardiac T-cell infiltration promoting diastolic dysfunction and hypertrophy.^4^ Guideline-directed medical therapy for heart failure with reduced ejection fraction (HFrEF) is of limited efficacy in HFpEF. Although sodium–glucose cotransporter 2 inhibitors and glucagon-like peptide-1/glucose-dependent insulinotropic polypeptide receptor agonists that reduce obesity can improve mortality and/or hospitalizations, cardiometabolic HFpEF remains a syndrome of great unmet clinical need.^5^

AKAP6β (mAKAPβ) is a 230 kDa scaffold protein present on the cardiomyocyte outer nuclear membrane that organizes a large multi-protein complex (“signalosome”) composed of signaling enzymes such as protein kinases and phosphatases and gene regulatory proteins such as transcription factors and histone acetylases.^6,7^ By integrating signaling by different second messenger pathways, AKAP6β signalosomes regulate stress-induced gene expression responsible for pathological cardiac remodeling.^6–8^ We have shown that targeting of AKAP6β signalosomes in mice can inhibit the development of heart failure due to chronic pressure overload.^8,9^ This included both AKAP6β gene knockout and treatment of mice with an AAV vector that expresses a competitive binding peptide comprising AKAP6β amino acid residues 1694-1833,^9^ herein called “Kinase-Binding Domain” (KBD), that was originally defined for its binding of p90 ribosomal S6 kinase type 3 (RSK3).^10^ Based upon these findings, we hypothesized that targeting of AKAP6β signalosomes would also be beneficial in HFpEF.

Here, we report that AKAP6β KBD peptide expression using a cardiotropic serotype 9 self-complementary AAV vector (AAV9sc.KBD,^11^ Figure S1) improved diastolic dysfunction and heart failure in a female swine model of cardiometabolic syndrome and HFpEF.^2,12^ Notably, T-cell infiltrates were reduced by the cardiomyocyte-directed therapy. Based upon an unbiased screen for new pro-inflammatory KBD binding partners whose binding to endogenous AKAP6β scaffold protein might be inhibited by AAV-expressed KBD peptide, evidence is provided that the KBD peptide binds CaMKII and that AKAP6β-bound CaMKII activates the NF-κB transcription factor pathway.^13–15^ A model is proposed that in response to antecedent “outside-in” inflammation in HFpEF, AKAP6β-CaMKII-dependent NF-κB activation in cardiomyocytes promoting inflammation comprises a positive feedback loop required for disease progression in cardiometabolic HFpEF.

## Methods

### Ossabaw Model of Cardiometabolic HFpEF

The Ossabaw swine strain has baseline metabolic abnormalities including insulin resistance.^16^ We have shown that when fed a high calorie, high fat diet, Ossabaw develop morbid obesity and metabolic syndrome, which in the presence of increased cardiac afterload progresses to a HFpEF-like syndrome.^12^ At two months of age, intact female Ossabaw were placed on a high fat diet, which continued for the duration of the 10 month study period (Figure 1A). At 6 months of age, before aortic banding to increase cardiac afterload, the pigs were tested for neutralizing antibodies and divided into two heart failure (HF) cohorts. Prior to banding, the two cohorts had similar body weights, insulin resistance, hypercholesterolemia, other serum chemistries, and cardiac function (Figures S2 and S3, Week 0). However, swine with lower AAV9 neutralizing antibody titers were preferred for AAV9sc.KBD treatment (Table S1). Immediately following banding (Figure S4), 7 swine were treated by peripheral intravenous infusion with 8 x 10^12^ viral genomes (vg) per kg body weight AAV9sc.KBD, and 5 swine with control saline infusion. Five additional 1-year-old Ossabaw swine raised on a standard normal chow were acquired as reference controls for endpoint studies.

**Figure 1.**
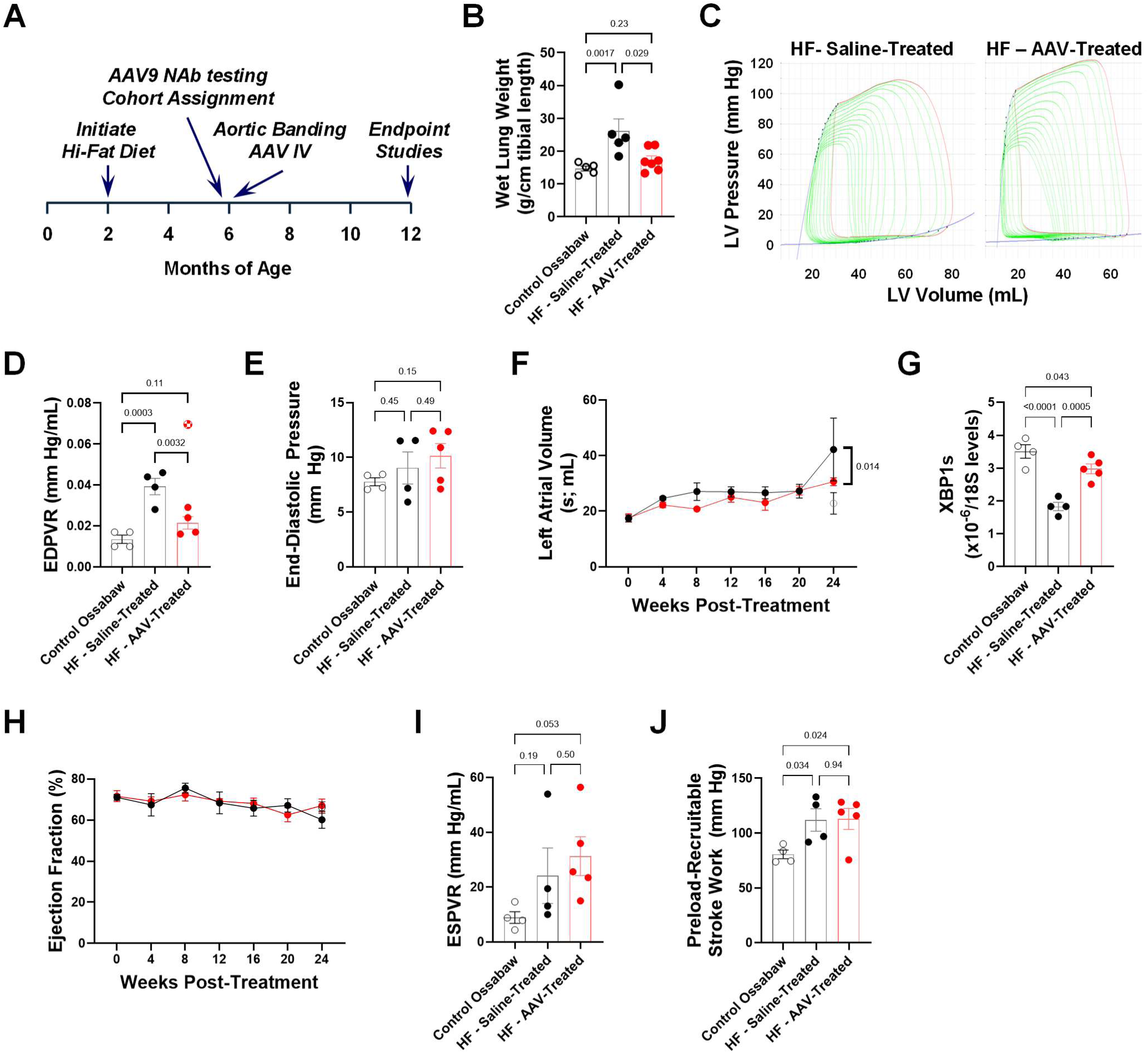
Prevention of diastolic dysfunction and heart failure in a swine model of cardiometabolic syndrome by AAV9sc.KBD gene therapy. **A.** Time-line for study of intact female Ossabaw swine. Two heart failure (HF) cohorts were fed a high-fat, high calorie diet starting at 2 months of age and subjected to aortic banding at 6 months of age, before being treated immediately after banding with 8 x 10^12^ viral genomes (vg) per kg body weight (3 x 10^14^ vg) AAV9sc.KBD (*n* = 7) or saline vehicle (Dulbecco’s Phosphate Buffered Saline with 0.001% Pluronic F68; *n* = 5) via peripheral intravenous infusion (Table S1). An additional control cohort included 1-year-old swine raised on a standard diet and studied only at endpoint (*n* = 5). *n* refers in this figure to the number of individual pigs. “Weeks Post-Treatment” in subsequent panels refers to time after aortic banding and AAV infusion. B. Wet lung weight at endpoint. n = 5,5,7 for control, HF – saline-treated and HF – AAV-treated Ossabaw, respectively. Data analyzed by Kruskal-Wallis test and Uncorrected Dunn’s tests. C. Representative pressure-volume loops at endpoint shown for saline-treated (pig 3367) and AAV-treated (pig 3340) HF swine. D. EDPVR (End-Diastolic Pressure Volume Relationship). n = 4 per cohort. AAV-treated pig 3391 (hatched red symbol) was an outlier (Grubbs α = 0.05) and excluded from EDPVR analysis. E. End-diastolic pressure at endpoint. F. Left atrial volume measured by serial echocardiography. G. XBP1s mRNA levels normalized to 18S RNA. H. Ejection fraction by serial echocardiography showed no significant difference between cohorts or between initial and end-points for the HF cohorts. I, J. ESPVR (End-Systolic Pressure Volume Relationship) and Preload-Recruitable Stroke work measured by pressure-volume loops. Panels D,E,G,I,J: Data analyzed by 1-way ANOVA and Uncorrected Fisher’s LSD tests. Panels E,G,I,J: n = 4,4,5 for control, saline- and AAV-treated HF Ossabaw, respectively. Panels F,H: Data for HF cohorts analyzed by matched (by subject) 2-way ANOVA (treatment and time) with mixed-effects analysis and Tukey’s test. n = 5 for control and HF – saline-treated Ossabaw, and 7 (0-20 weeks) or 5 (24 weeks, see Table S1) for HF-AAV-treated swine.

### Other Methods and Large Datasets

Detailed methods are provided in the online Supplemental Material. Single nucleus and bulk RNA-sequencing data (snRNA-seq and RNA-seq) are available at NCBI GEO GSE335365 and GSE335057, respectively. Raw peptide and search results files for immunoprecipitation-mass spectrometry are available at https://massive.ucsd.edu. dataset ID: MSV000101850. Most data supporting the conclusions of this article are included within the article and Supplemental Material. Other data and materials used during the study are available from the corresponding authors on reasonable request.

## Results

### Treatment of a swine model of cardiometabolic HFpEF with a AKAP6β-derived gene therapy

The combination of high fat diet and aortic banding resulted in heart failure in female Ossabaw swine,^12^ demonstrated by increased wet lung weight in the saline-treated HF cohort at endpoint (Figure 1B). Notably, AAV9sc.KBD treatment prevented the increase in wet lung weight. The beneficial effect of gene therapy in the HFpEF model was associated with improved diastolic function. Terminal LV pressure-volume loops showed that end-diastolic pressure-volume relationship (EDPVR) was increased for saline-treated HF swine 2.9-fold compared to control Ossabaw, whereas there was no significant increase in EDPVR for the AAV-treated HF cohort (Figure 1C,D). Left ventricular end-diastolic pressure was the same as control for both HF cohorts (Figure 1E), consistent with observations that end-diastolic pressure can be normal in HFpEF patients at rest.^17^ Corroborating the EDPVR results, left atrial volume was significantly less in the AAV-treated than saline-treated HF cohort at endpoint (Figure 1F). Decreased expression of the active spliced form of the X-box binding protein 1 transcription factor (XBP1s) is a HFpEF-specific marker in human patients and promotes HFpEF in mice.^18,19^ In comparison to control Ossabaw, LV XBP1s mRNA was significantly decreased in AAV-treated compared to saline-treated HF (Figure 1G and Figure S5).

Cardiac remodeling of the HF cohorts as indicated by increased LV posterior wall thickness, relative wall thickness, and calculated LV mass tended to be less for AAV9sc.KBD-treated swine at later timepoints (Figure S3E,H,I), albeit overall cardiac hypertrophy at endpoint was not reduced by AAV9sc.KBD treatment (Table S1). Importantly, systolic function was not decreased by the gene therapy. Ejection fraction was preserved for all cohorts throughout the study period (Figure 1H), and elevated contractility in both HF cohorts (end-systolic pressure-volume relationship and preload-recruitable stroke work) compared to control Ossabaw was not reduced (Figure II,J). Taken together, AAV9sc.KBD treatment prevented the development of diastolic dysfunction and heart failure in a swine model of cardiometabolic HFpEF without adverse effects on the heart.

While improving cardiac function, the cardiomyocyte-specific gene therapy did not affect the extent of metabolic syndrome. At endpoint, the HF cohorts were ∼70% greater in weight than the control Ossabaw (Table S1 and Figure S3A, Week 24). The HF cohorts continued to have hypercholesterolemia whereas all three cohorts exhibited insulin resistance, as expected^16^ (Figure S6). As in our previous study,^11^ intravenous administration resulted in substantial AAV9sc.KBD delivery in all organs tested (Table S2). Importantly, KBD transcripts were selectively expressed in the cardiac left and right ventricles, including septum, with minimal detection in liver and skeletal muscle (SKM). KBD mRNA left ventricle (LV) expression was confirmed by RNAscope *in situ* hybridization, with detection of mature cytosolic transcripts in LV myocardium only with an antisense but not sense control probe (Figure S7 and Table S3).

### Inhibition of cardiac immune signatures and T-cell infiltration in KBD-treated swine

To study the mechanisms conferring KBD peptide benefit, swine LV tissue acquired at endpoint was analyzed by bulk RNA sequencing (RNA-seq). 250 genes were differentially expressed between the saline-treated HF cohort and control Ossabaw (Table S4, adjusted p-value < 0.05). Heatmap analysis and comparison of the log_2_-fold changes in expression of these genes due to HF induction (saline-treated HF cohort vs. control Ossabaw) versus following KBD peptide expression (AAV-treated vs. saline-treated HF cohort) showed that AAV9sc.KBD treatment partially reversed the HF-associated gene expression program (Figure 2A,B). For HF-associated genes, immune stress responses and cell signaling dominated the list of gene ontologies (Figure 2C and Table S5). Additionally, Ingenuity Pathway Analysis showed increased gene expression related to interferon α/β (p = 4.3e-17) and interferon γ (p = 2.0e-11) signaling pathways, like previously reported findings in a mouse HFpEF model.^4^ Interestingly, of the 250 genes, 109 have been reported to be regulated directly by the pro-inflammatory transcription factor Nuclear Factor κB (NF-κB, Table S4).

**Figure 2.**
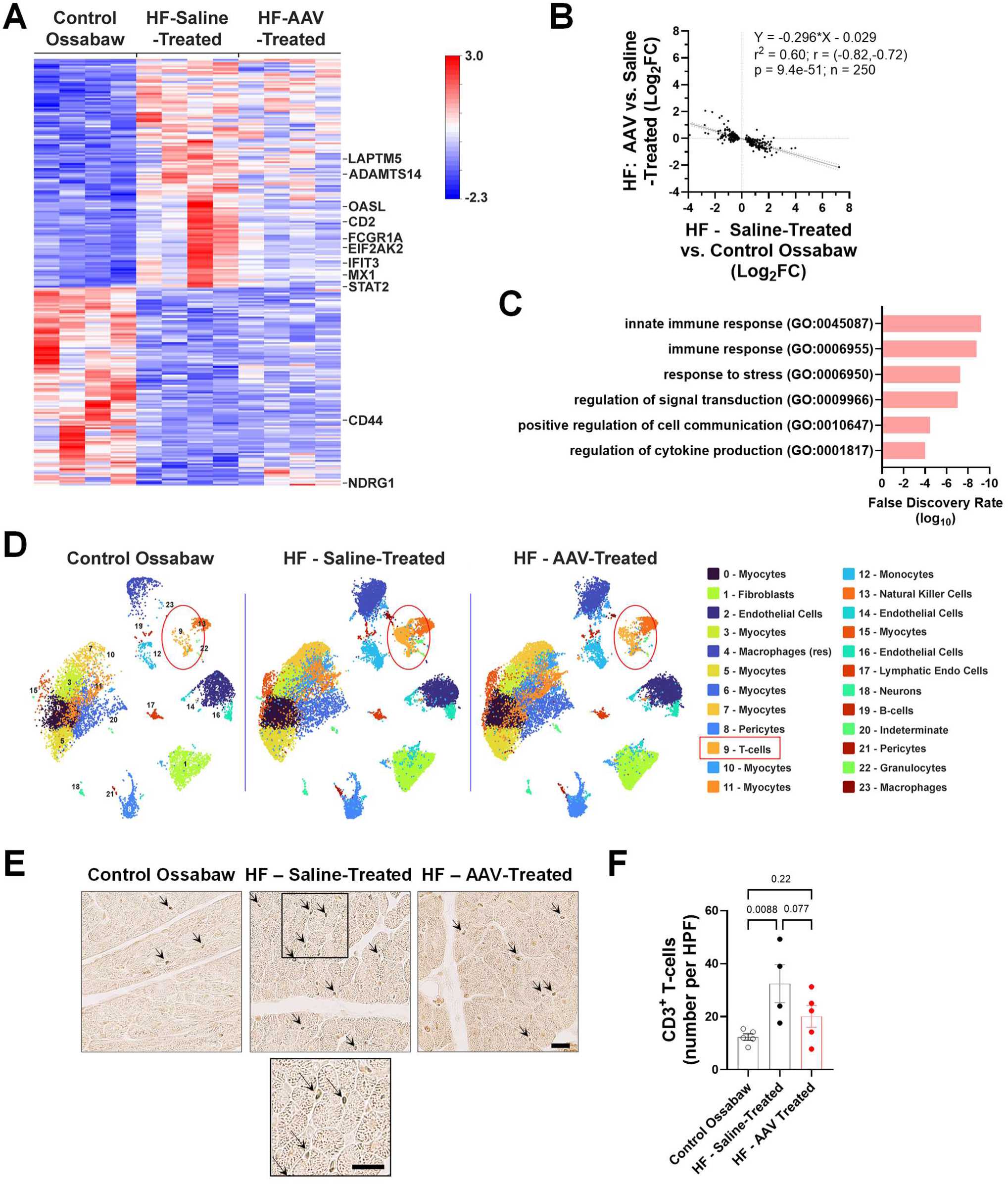
AAV9sc.KBD inhibits T-cell Inflammation in the swine cardiometabolic syndrome model. **A.** Heatmap of the 250 genes differentially expressed in saline-treated HF and control Ossabaw swine as determined by bulk RNA-seq for LV tissue (n = 4 per cohort, adjusted *p* < 0.05, z-score scale shown). See Table S4. **B.** Linear regression showing inverse relationship between the effects (log_2_ fold change [Log_2_FC]) of disease (X-axis: saline-treated HF vs. control Ossabaw swine) and AAV9sc.KBD treatment (Y-axis: AAV-treated vs. saline-treated HF Ossabaw) for the genes shown in A. **C.** Selected gene ontologies for the gene set in A. See Table S5 for additional gene ontologies. **D.** Uniform Manifold Approximation and Projection (UMAP) for single-nuclear RNA-seq (snRNA-seq) analysis of LV tissue by cohort with clusters annotated by cell type. T-cell cluster 9 circled red. See Figure S8 and Table S6. n = 3,4,4 for control, HF – saline-treated and HF – AAV-treated Ossabaw, respectively. **E.** Staining of swine tissue for the pan-T-cell marker CD3. Scale bars – 20 µm (large sections at same scale). **F.** Quantification of CD3 staining in E. n = 5,4,5 for control, HF – saline-treated and HF – AAV-treated Ossabaw, respectively. Data analyzed by 1-way ANOVA and Uncorrected Fisher’s LSD tests. *n* refers in this figure to the number of individual pigs.

Given the RNA-seq results and the role of inflammation in HFpEF,^3^ single nucleus RNA-seq (snRNA-seq) analysis was used to identify immune cell infiltrates in swine LV myocardium (Figure 2D). 24 clusters distinguished by Uniform Manifold Approximation and Projection (UMAP) were identified by canonical cell markers (Figure S8A,B and Table S6). As expected, AAV9sc.KBD mRNA was detected only in the AAV-treated cohort and >99.4% in cardiomyocytes, which were represented by 8 clusters comprising 51% of the sequenced nuclei (Figure S8C). Comparison of cluster nuclei counts showed that T-cells were increased in the saline-treated HF cohorts compared to control Ossabaw and AAV-treated cohorts (Figure 2D and Figure S8D). The increased T-cell infiltrates in the saline-treated HF pigs compared to the other cohorts was corroborated by immunohistochemical staining for CD3 antigen showing intramyocardial T-cells (Figure 2E,F).

In further snRNA-seq analysis, CD45^+^ (PTPRC-expressing) immune cell nuclei were segregated into 17 distinct clusters (Figure S9A,B and Table S7). T-cells (naïve, mature, double negative and γ/δ represented by 5 clusters) were over-represented in the saline-treated HF cohort 5-8-fold compared with control Ossabaw and AAV-treated HF cohorts (p < 0.005). Notably, mature T-cells comprising cluster 0, including both CD4^+^ and CD8^+^ cells, were enriched 8-15-fold in the saline-treated HF cohort compared to the other cohorts (p < 0.0001, Figure S9C). Consistent with the above gene ontology analysis (Figure 2C), HFpEF involves both adaptive and innate immune responses.^4,20^ Resident macrophages tended to be differentially clustered between the control Ossabaw and HF cohorts but were not affected by the AAV gene therapy (Figure S9C, compare clusters 2 and 4). Taken together, these results suggest AAV9sc.KBD treatment selectively reduces T-cell inflammation prominent in cardiometabolic syndrome.

### Identification of CaMKII as an AKAP6β-binding partner

The AKAP6β KBD was initially defined by its RSK3 binding.^10^ However, RSK3 is not associated with inflammation. To identify other AKAP6β KBD protein binding partners whose AKAP6β binding would be inhibited by the AAV-expressed recombinant KBD peptide and explain the observed inhibition of T-cell inflammation, adult rat ventricular myocytes expressing epitope-tagged KBD peptide were used for antibody immunoprecipitation-mass spectrometry analysis (Figure 3A and Table S8). Besides the KBD peptide, tryptic peptides unambiguously identified CaMKII δ, γ, and α (in decreasing order of detection). KBD-CaMKII binding was validated by co-expression of recombinant KBD peptide and CaMKII proteins in COS-7 cells (Figure S10 A,B). CaMKII δC and γX1 alternatively-spliced *Camk2d* and *Camk2g* isoforms^21^ both co-immunoprecipitated with KBD. In addition, consistent with the KBD peptide comprising a native AKAP6β domain, endogenous AKAP6β and CaMKII were co-immunoprecipitated with anti-CaMKII antibody from neonatal rat ventricular myocyte extracts (Figure 3B). Importantly, KBD peptide expression inhibited AKAP6β-CaMKII co-immunoprecipitation (Figure 3B,C), defining the KBD as an anchoring disruptor peptide for not only RSK3, but also CaMKII. We considered that the RSK3 and CaMKII KBD binding sites might be independent. However, mapping studies showed that the kinases bind overlapping KBD peptide fragments, such that we were unable to separate the RSK3 and CaMKII binding sites (Figure S10B).

**Figure 3.**
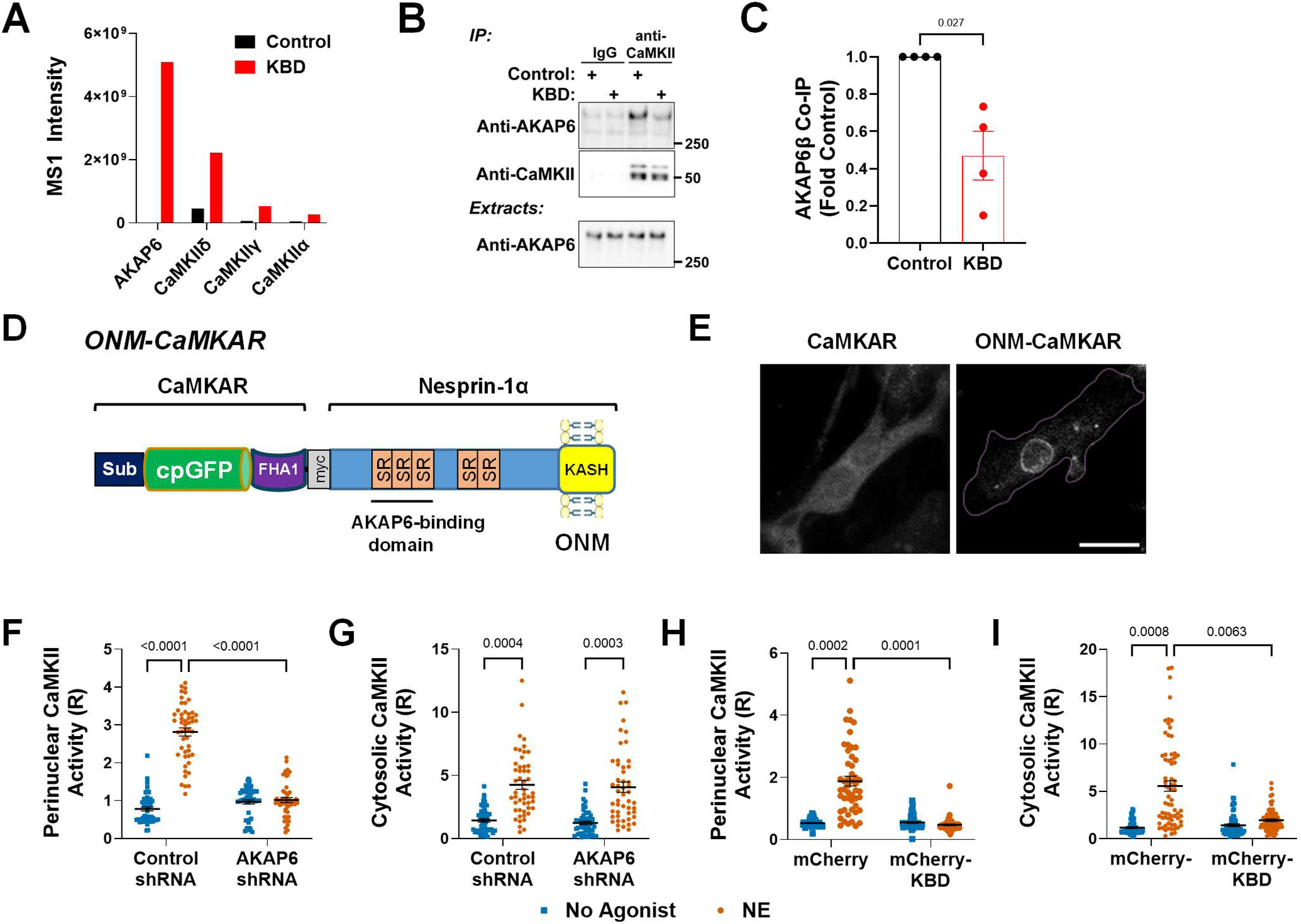
Ca^2+^/calmodulin-dependent protein kinase II binds the Kinase-Binding Domain (KBD) of AKAP6β. **A.** To identify novel KBD binding partners in an unbiased screen, adult rat ventricular myocytes expressing myc- (and GFP-) tagged recombinant KBD peptide were used to prepare cross-linked extracts for myc-tag antibody immunoprecipitation and mass spectrometry. Control samples were not transduced with KBD adenovirus. The sum of MS1 intensities for peptides identified for the indicated proteins are provided as a measure of protein detection. See Table S8 for peptide details. AKAP6 peptides were all from the KBD. **B, C.** Protein complexes were immunoprecipitated using CaMKIIδ antibody from neonatal rat ventricular myocytes expressing either GFP-tagged KBD or GFP control. AKAP6β was detected by western blot in total extracts (input) and immunoprecipitates (IP). n = 4 independent experiments, compared by paired t-test. **D.** Design of an outer nuclear membrane-targeted CaMKII activity reporter (ONM-CaMKAR). CaMKAR ratiometric sensor contains circularly permuted green fluorescent protein (cpGFP) fused on the N- and C-terminus to CaMKIIα aa 281-291 peptide containing the Thr-286 autophosphorylation site (Sub) and to a FHA1 phospho-peptide binding domain, respectively.^22^ The ratio of emission at 520 nm (R) due to excitation at 488 versus excitation at 405 nm indicates relative CaMKII activity. Nesprin-1α contains 3 spectrin-like repeats (SR) and the ONM-specific Klarsicht, ANC-1, Syne homology (KASH) transmembrane domain.^23^ **E.** CaMKAR and ONM-CaMKAR fluorescence in neonatal rat ventricular myocytes. Scale bar – 20 µm. Cell in right panel is outlined in purple. **F-I**. Neonatal myocytes expressing either ONM-CaMKAR (F,H) or parent CaMKAR (G,I) biosensor and either AKAP6 or control shRNA (F,G) or mCherry-tagged KBD or mCherry control (H,I) were cultured overnight in the presence of norepinephrine (NE, 10 nmol/L) or no agonist before ratiometric imaging. R is ratio of emission intensity for 488 versus 405 nm excitation. Parental CaMKAR signal was quantified only within the cytosol. 50-70 myocytes were imaged across 4 biological replicates per condition. Data were analyzed by nested ANOVA and uncorrected Fisher’s LSD tests.

The interaction of CaMKII with AKAP6β and the KBD peptide was studied further in live cells using the ratiometric fluorescent CaMKII activity reporter CaMKAR.^22^ To compare cytosolic CaMKII activity detected with the parent CaMKAR sensor with CaMKII activity in the perinuclear AKAP6β compartment, CaMKAR was expressed in fusion to nesprin-1α, the outer nuclear membrane protein responsible for AKAP6β localization in myocytes (ONM-CaMKAR, Figure 3D,E).^23,24^ Consistent with previous findings that active CaMKIIδC accumulates at the nuclear envelope in stressed cardiomyocytes,^25^ ONM-CaMKAR signal was detected in neonatal myocytes after prolonged Ca^2+^ ionophore stimulation (ionomycin, Figure S10C). Likewise, CaMKII activity was detected by both ONM-CaMKAR and cytosolic parental CaMKAR sensors in myocytes treated overnight with norepinephrine (NE, Figure 3F-I). Consistent with AKAP6β being a scaffold for perinuclear CaMKII, AKAP6β expression was required for CaMKII activity detected with ONM-CaMKAR, but not cytosolic CaMKAR (Figure 3F,G). In addition, consistent with inhibition of AKAP6β-CaMKII binding, expression of a mCherry-KBD fusion protein inhibited perinuclear CaMKII activity (Figure 3H). KBD peptide expression also inhibited cytosolic CaMKII activity (Figure 3I), suggesting that KBD peptide binding can inhibit CaMKII catalytic activity independently of blocking AKAP6β binding. However, in all *in vitro* experiments, recombinant KBD peptide and control proteins were expressed at much higher levels than the non-tagged KBD peptide in swine (data not shown).

### AKAP6β-CaMKII signalosomes regulate the pro-inflammatory NF-κB pathway

In cardiomyocytes, CaMKII phosphorylates proteins regulating excitation-contraction coupling, gene expression, and other cellular processes.^13,14^ To screen for CaMKII substrates whose phosphorylation was inhibited by KBD peptide expression in the Ossabaw swine model, LV extracts were analyzed by western blotting (Figure 4). Notably, phosphorylation of the CaMKII site Ser-177 on Inhibitor of NF-κB (IκB) kinase (IKK) was increased 3-4-fold in saline-treated HF swine when compared to control Ossabaw, but not in AAV9sc.KBD-treated HF swine (Figure 4A,B). In contrast, the CaMKII substrates endothelial nitric oxide synthase (eNOS Ser-1177),^26^ cAMP Response Element-Binding protein (CREB Ser-133),^27^ and PLN Thr-17 were not significantly increased in phosphorylation in the Ossabaw model, whereas histone deacetylase 4 (HDAC4 Ser-632)^28^ was decreased in phosphorylation (Figure 4A,C).^14^ In addition, PLN Ser-16 (PKA site) and ERK1/2 (activation sites Thr-202/Tyr-204) were not differentially phosphorylated among the three cohorts. The RSK3 substrate SRF Ser-103 was decreased in phosphorylation in the HF cohorts at endpoint.

**Figure 4.**
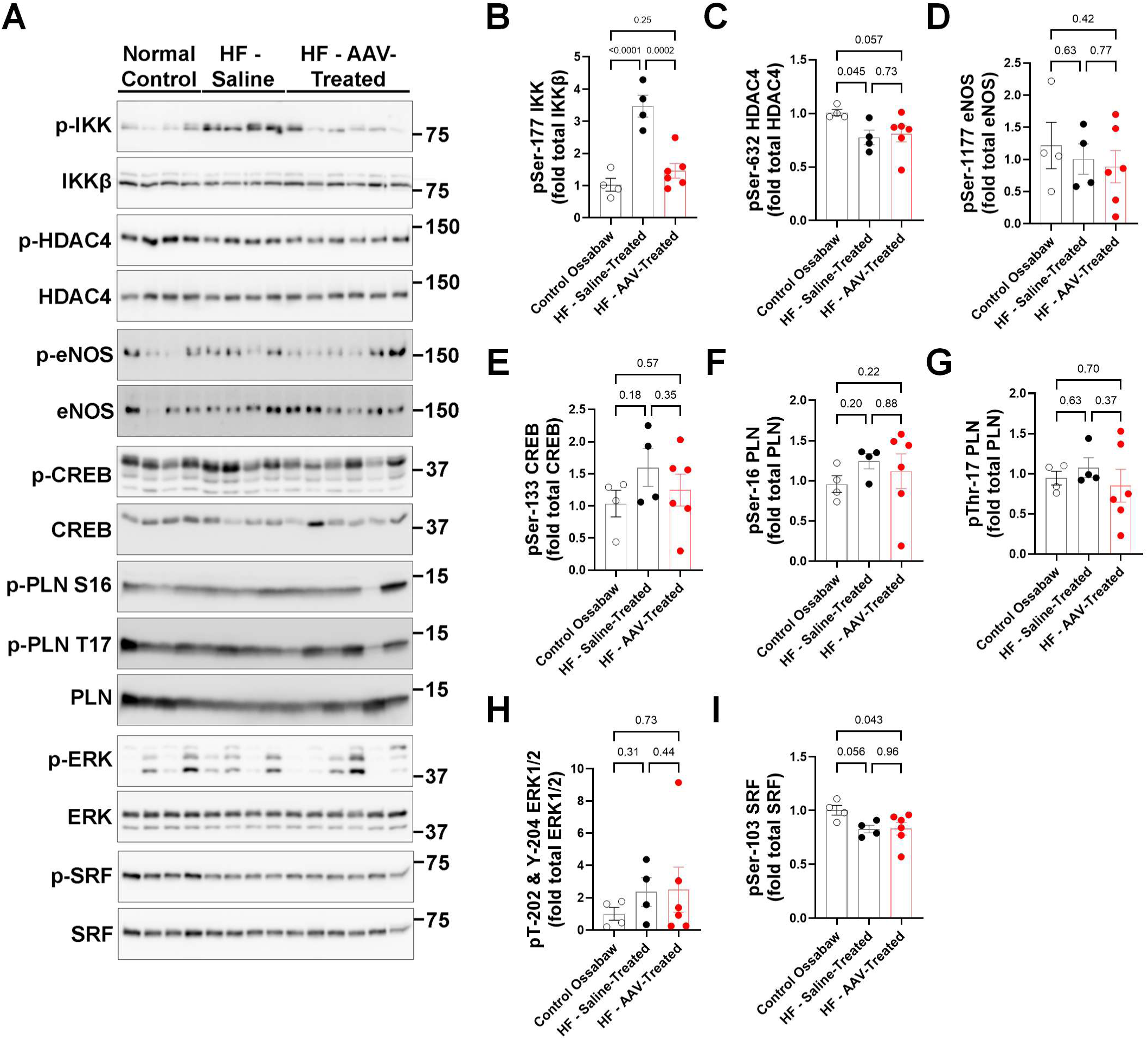
Regulation of signaling pathways in the Ossabaw cardiometabolic syndrome model. **A.** Left ventricular tissue (or in the case of HDAC4, HDAC4 antibody immunoprecipitates) was analyzed by western blotting for the indicated phospho-proteins, with data normalized by the corresponding total protein levels: **B.** Inhibitor of NF-κB Kinase Subunit (IKK) phospho-serine 177; **C.** histone deacetylase 4 (HDAC4) phospho-serine 632; **D.** endothelial nitric oxide synthase (eNOS) phospho-serine 1177; **E.** cAMP Response Element Binding Protein (CREB) phospho-serine 133; **F.** phospholamban (PLN) phospho-serine 16; **G.** PLN phospho-threonine 17; **H.** extracellular signal-regulated kinase 1/2 (ERK1/2) phospho-threonine 202 and phospho-tyrosine 204; **I.** serum response factor (SRF) phospho-serine 103. All data were analyzed by 1-way ANOVA and Uncorrected Fisher’s LSD tests, except for F and H which were analyzed by Kruskal-Wallis and uncorrected Dunn’s tests. n = 4,4,6 for control, HF – saline-treated and HF – AAV-treated Ossabaw, respectively. *n* refers to the number of individual pigs.

NF-κB transcription factor regulates pro-inflammatory gene expression.^29^ In unstimulated cells, NF-κB is retained in the cytoplasm by IκB protein. Canonical NF-κB activation involves the activation of IKK complex composed of IKK α, β, and γ subunits, which phosphorylates IκB inducing IκB ubiquitination and degradation and NF-κB nuclear translocation. CaMKII phosphorylates IKKβ and IκBα, driving NF-κB-dependent inflammation.^15,30–32^ As KBD peptide expression inhibited IKK phosphorylation in swine, we hypothesized that IKKβ associates with cardiomyocyte AKAP6β signalosomes where IKKβ is phosphorylated by AKAP6β-dependent CaMKII in cardiometabolic HFpEF. First, we showed by co-immunoprecipitation assay that IKKβ bound AKAP6β when expressed in COS-7 cells (Figure 5A). Next, KBD peptide, which competes AKAP6β-CaMKII binding, was shown to inhibit NE-induced IKK phosphorylation in adult rat cardiomyocytes (Figure 5B). In neonatal rat cardiomyocytes, NE-induced NF-κB nuclear translocation depended upon both CaMKII activity and AKAP6β expression and was inhibited by KBD peptide expression (Figure 5C-F). Likewise, NE-induced NF-κB nuclear translocation was dependent upon AKAP6β expression in adult rat cardiomyocytes (Figure 5G,H). Taken together, these results suggest that in cardiomyocytes AKAP6β-bound CaMKII phosphorylates IKKβ, driving NF-κB nuclear translocation.

**Figure 5.**
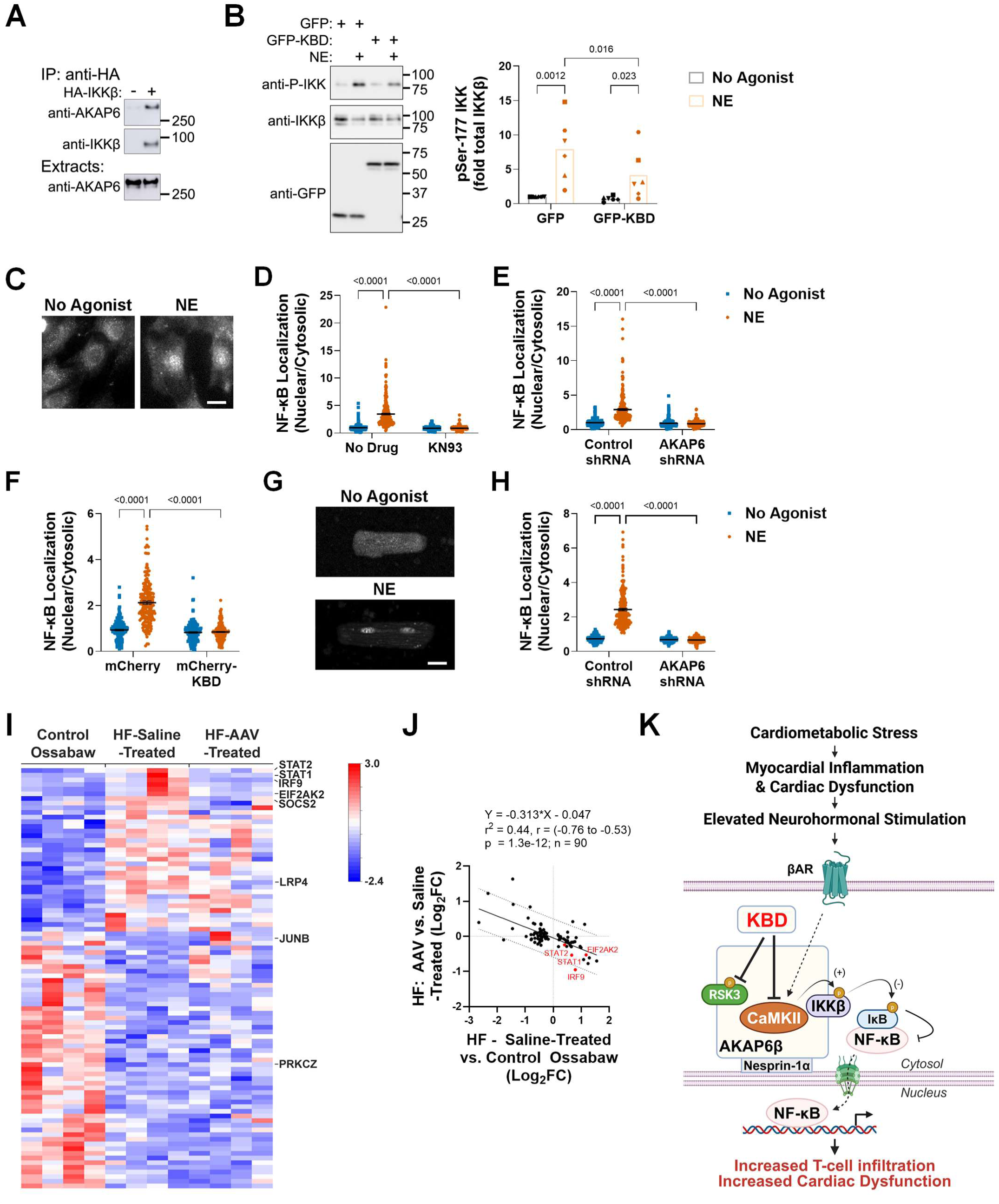
Regulation of NF-κB by AKAP6β-CaMKII signalosomes in cardiomyocytes. **A.** Protein complexes in COS-7 cells expressing HA-tagged IKKβ and (myc-tagged) AKAP6β were immunoprecipitated using HA antibody and detected by western blotting with AKAP6 and IKKβ antibodies. n = 3 independent experiments. **B.** Adult rat ventricular myocytes expressing GFP or GFP-tagged KBD were treated for 30 minutes with norepinephrine (NE, 10 µmol/L) before whole cell extracts were analyzed by western blotting with the indicated antibodies. n = 6 independent myocyte preparations indicated by different symbols. Data analyzed by matched (by biological replicates) 2-way ANOVA and uncorrected Fisher’s LSD tests. **C.** Neonatal rat ventricular myocytes were cultured in minimal medium with and without 10 nmol/L NE for 2 days before staining with a p65 NF-κB antibody and imaging by widefield fluorescence microscopy for fluorescence intensity in the cytosol and nucleus. Scale bar – 20 µm. **D.** Neonatal myocytes were treated as in C except for the inclusion of the CaMKII inhibitor KN-93 (5 µmol/L) as indicated. **E.** Neonatal myocytes transduced with adenovirus expressing control or AKAP6 shRNA were cultured and analyzed as in C. **F.** Neonatal myocytes transfected with expression plasmids for mCherry-KBD fusion protein or mCherry were cultured and analyzed as in C. **G.** Adult rat ventricular myocytes were cultured in minimal medium with and without 10 nmol/L NE for 24 hours before staining with a p65 NF-κB antibody and imaging by confocal fluorescence microscopy. Scale bar – 20 µm. **H.** Adult myocytes transduced with adenovirus expressing control or AKAP6 shRNA were cultured and analyzed as in G. D-F, H. Mean nuclear to cytosolic ratio for NF-kB fluorescence is shown for 50-85 cells per biological replicate. n = 4,4,3,4 independent myocyte preparations for D,E,F,H, respectively. Data analyzed by nested 1-way ANOVA and Uncorrected Fisher’s LSD tests. **I.** Heatmap of the 90 NF-κB-regulated genes differentially expressed in cardiomyocytes between saline-treated heart failure and control Ossabaw cohorts as defined by BayesPrism analysis of snRNA-seq data (p-value < 0.05, z-scores shown). See Table S9 for additional information. **J.** Linear regression showing inverse relationship between the effects (log_2_ fold change [Log_2_FC]) of disease (X-axis: saline-treated HF vs. control Ossabaw swine) and AAV9sc.KBD treatment (Y-axis: AAV-treated vs. saline-treated HF Ossabaw) for the genes shown in I. **K.** Model for regulation of myocardial inflammation in cardiometabolic HFpEF by AKAP6β signalosomes and its inhibition by AAV9sc.KBD gene therapy. CaMKII induces IKKβ-dependent activation of NF-κB transcription factor at the perinuclear AKAP6β signalosome, exacerbating T-cell infiltration and inflammation induced by metabolic syndrome and systemic inflammation. Expression of the KBD peptide comprising the AKAP6β binding domain for CaMKII inhibits pro-inflammatory signaling. The KBD peptide can also inhibit RSK3-dependent pathological remodeling. Created in BioRender. Kapiloff, M. (2026) https://BioRender.com/a0s0esa

To obtain evidence that NF-κB-dependent cardiomyocyte gene expression was relevant to the effects of AAV9sc.KBD in the cardiometabolic Ossabaw swine model, we turned again to the RNA-seq datasets. BayesPrism analysis is a method for the deconvolution of RNA-seq datasets to determine cell-type specific gene expression when scRNA- or snRNA-seq data are available.^33^ Of the cardiomyocyte-expressed genes, 363 were differentially expressed between saline-treated HF and control Ossabaw cohorts (Table S9). Like the parent RNA-seq datasets, heatmap analysis and comparison of the log_2_-fold changes in expression of these genes due to HF induction (saline-treated HF cohort vs. control Ossabaw) versus following KBD peptide expression (AAV-treated vs. saline-treated HF cohort) showed that AAV9sc.KBD treatment partially reversed the HF-associated gene expression program (Figure S11). Of the 363 differentially-expressed genes, 90 have been reported to be NF-κB regulated (Table S9), 35 being activated and 55 repressed. Heatmap analysis and comparison of log_2_-fold changes similarly showed a partial reversion of pathological gene expression following AAV treatment (Figure 5I,J). Taken together, these results suggest that suppression of the NF-κB-dependent pro-inflammatory transcriptomic program in cardiomyocytes contributes to the KBD gene therapy-dependent improvement in cardiac status in the Ossabaw swine model of cardiometabolic HFpEF.

## Discussion

Proof-of-concept is provided here that AAV9sc.KBD gene therapy can inhibit the development of diastolic dysfunction, myocardial inflammation, and heart failure in a female Ossabaw swine model of cardiometabolic HFpEF.^2,12^ Mechanistic insight is provided through RNA-seq and snRNA-seq analyses that cellular and molecular hallmarks of HFpEF including T-cell infiltration and NF-κB-dependent gene expression were attenuated by AAV9sc.KBD therapy. CaMKII and IKKβ were shown to bind the AKAP6β scaffold protein, and AKAP6β-CaMKII signalosomes to activate NF-κB in cardiomyocytes.

Functionally, a significant outcome of the study was that AAV9sc.KBD was able to prevent the development of heart failure. The gene therapy prevented the devastating link often observed in HFpEF (and Ossabaw swine with cardiometabolic HFpEF) by which diastolic dysfunction leads to atrial remodeling and lung congestion via elevated filling pressures realized either at rest or during stress (e.g., exercise or activities of daily living). Indeed, AAV-treatment prevented pathological increases in EDPVR that were associated with normal wet lung weight and attenuated left atrial remodeling. Further, AAV9sc.KBD did not impair systolic function and was related to trends towards decreased pathological LV remodeling.

Mechanistically, the most provocative finding in this study was the diminution of T-cell infiltrates in AAV-treated HF swine. Whereas inflammation in HFrEF can be described as “inside-out” because it is induced by myocardial damage, HFpEF is associated with elevated systemic inflammation that negatively affects the heart, i.e. “outside-in.”^3^ For example, cardiometabolic syndrome is associated with elevated blood cytokine and T-cell levels consistent with activation of adaptive immunity. As found here in Ossabaw swine, we have shown in mice that a HFpEF-like phenotype is associated with elevated T-cell (both CD4^+^ and CD8^+^) and monocyte populations in the myocardium, with downregulation in T-cells of XBP1s expression and the unfolded protein response pathway.^4^ Importantly, T-cell deficient *Tcra^-/-^* mice exhibited decreased cardiac hypertrophy and diastolic dysfunction in response to high fat-diet and L-NAME administration.^4^ Here, a cardiomyocyte-directed gene therapy inhibited myocardial T-cell infiltration. This suggests that T-cells and cardiomyocytes form a positive feedback loop, in which inflammation-induced pathological remodeling results in pro-inflammatory cardiomyocyte signaling that potentiates the local immune reaction (Figure 5K). AAV9sc.KBD therapy apparently disrupts this positive feedback by inhibiting cardiomyocyte signaling important for enhancing T-cell infiltration, proliferation, or survival within myocardium. Thus, this study suggests that KBD peptide therapy inhibits pathological cardiac remodeling through both inhibition of myocyte autonomous mechanisms and secondarily to decreased inflammation.

Consistent with the prominent role of T-cells and both adaptive and innate immunity in HFpEF and as previously reported in this Ossabaw HFpEF model,^4,12,20^ transcriptome analysis showed that pathways for the innate immune response and cytokine production were activated in HF swine, including NF-κB-regulated genes. The expression of some NF-κB-regulated genes was normalized by AAV9sc.KBD treatment, for example, eukaryotic translation initiation factor 2-alpha kinase 2 (*EIF2AK2*), interferon regulatory factor 9 (*IRF9*), and signal transducer and activator of transcription 1 and 2 (*STAT1* and *STAT2*) expressed in cardiomyocytes. The inflammasome is important for post-translational processing of cytokines that promote myocardial inflammation driving fibrosis and cardiac dysfunction.^15^ EIF2AK2 binds the inflammasome protein NLR family pyrin domain containing 3 (NLRP3), inducing in cardiomyocytes interleukin-1β (IL-1β) proteolytic activation.^34^ IL-1β contributes to macrophage and T-cell myocardial recruitment.^15^ Independently of the inflammasome, IRF9, STAT1, and STAT2 form a transcription factor complex (interferon-stimulated gene factor 3) that in cardiomyocytes promotes cell death and myocardial inflammation.^35^

In an unbiased screen, CaMKII was identified as a new AKAP6β KBD binding partner. We detected AKAP6β-associated CaMKII activity only after prolonged Ca^2+^ elevation, consistent with previous findings that perinuclear active CaMKII is increased *in vivo* in response to chronic pressure overload.^25^ Whereas it was expected that the KBD peptide would competitively inhibit CaMKII binding to endogenous AKAP6β, thereby inhibiting perinuclear kinase activity, it was unexpected that the peptide would inhibit un-anchored CaMKII catalytic activity as detected using the cytosolic CaMKAR sensor. That KBD-binding inhibited CaMKII catalysis, whereas the complexing of CaMKII with AKAP6β was found necessary for compartmentalized perinuclear CaMKII activity, is paradoxical considering that KBD is a fragment of AKAP6β. The phosphatase calcineurin binds AKAP5 when inactive and is released upon activation to dephosphorylate nearby nuclear factor of activated T-cells (NFAT) transcription factor.^36^ AKAP6β might similarly serve as a sink for inactive CaMKII, which following activation dissociates from the scaffold and phosphorylates nearby substrates, like IKKβ. Alternatively, like AKAP7δ which regulates CaMKII-dependent phospholamban phosphorylation,^37^ AKAP6β might contain a second site that binds activated CaMKII. In that scenario, the KBD site serves as a sink for inactive CaMKII, whereas transfer of activated CaMKII to the second site promotes local substrate phosphorylation. Additional structure-function studies will be required to define the relevant mechanism.

Despite inhibiting IKK phosphorylation, cardiomyocyte-selective KBD peptide expression in swine did not inhibit CaMKII-dependent PLN Thr-17 phosphorylation, which is dependent upon a different AKAP (AKAP7δ) in cardiomyocytes.^37^ This contrasts with the marked suppression of PLN Thr-17 phosphorylation in transgenic mice expressing autocamtide-3 related inhibitory peptide (AiP), which is a competitive CaMKII inhibitor.^38^ Interestingly, KBD peptide expression had no deleterious effect on systolic function in the cardiometabolic swine model, whereas the AiP transgenic mice exhibited decreased systolic function during acute pressure overload.^38^ The differential effects of AAV9sc.KBD on IKK and PLN phosphorylation suggests that the relatively low level of KBD peptide expression in swine permitted selective inhibition of CaMKII compartments and is potentially indicative of the relative contributions of anchoring disruption and catalysis inhibition. Moreover, whereas NF-κB-dependent gene expression was regulated in the swine cardiomyocytes, neither PLN nor the CaMKII substrate HDAC4 was increased in phosphorylation. CaMKII phosphorylation of class IIA HDACs has been proposed to be regulated by inner nuclear membrane inositol-3-phosphate receptors,^28,39,40^ a compartment likely independent of AKAP6β at the outer nuclear membrane.^41,42^ These results underscore the importance of compartmentation to the intracellular signaling network, in which signaling pathways differentially regulate the activation of relevant downstream effectors in different intracellular compartments in response to different upstream stimuli and stress conditions.

The discovery that the AKAP6β KBD binds not only RSK3, but also CaMKII suggests that bivalency may result in the peptide’s broad efficacy in the prevention of heart failure. Cardiomyocyte RSK3 has been implicated in heart failure due to pressure overload, Hypertrophic Cardiomyopathy, and RASopathy and was previously observed to be elevated in aortic-banded Yucatan swine,^9,10,43,44^ suggesting inhibited RSK3-SRF signaling may have contributed to the trend towards improved cardiac remodeling in the AAV-treated Ossabaw HFpEF swine in the current study. However, consistent with observations suggesting that RSK3-SRF signaling comprises an early stress response,^9^ SRF phosphorylation was not increased at endpoint when LV tissue became available and was not investigated further herein. The multiple alternatively-spliced CaMKII isoforms expressed in cardiomyocytes within diverse intracellular compartments have been implicated in a wide range of physiological and pathological cardiac conditions. CaMKII contributes to the fine tuning of physiological excitation-contraction coupling, which is apparently essential in acute pressure overload.^38^ However, excessive CaMKII activity promotes arrhythmia, mitochondrial dysfunction, myocyte death due to ischemia/reperfusion injury, pressure overload-induced remodeling, diabetic cardiomyopathy, and, as discussed here, myocardial inflammation.^13,14,45^ AKAP6β gene deletion was beneficial in both mouse models of ischemic cardiomyopathy and chronic pressure overload,^7,8^ and KBD expression was found effective in mice subjected to pressure overload resulting in HFrEF and now here in a swine cardiometabolic HFpEF model.^9^ Although RSK3 global knockout is well tolerated in mice,^9,10^ CaMKII inhibition *in vivo* is complicated by the major role CaMKII serves in learning and memory, as well as cardiac physiology.^13^ The apparent selectivity of KBD peptide action and the possibility of cardiomyocyte-specific expression may confer advantages to a KBD gene therapy in the treatment of conditions involving CaMKII activation in the AKAP6β signalosome compartment, as will be investigated further in additional pre-clinical studies.

Limitations includes that due to the pigs’ weight at treatment and AAV cost, saline vehicle infusion was used instead of a control AAV. In addition, the AAV9sc.KBD dose was relatively low (8 x 10^12^ vg/kg).^11^ Both *in situ* hybridization and snRNA-seq detected KBD transcripts in some but not all cardiomyocytes, raising the possibility of a stronger effect size for higher doses. Local administration was not attempted given our previous findings that cardiac AAV9 delivery in swine is similar by intravenous and antegrade intracoronary coronary infusion.^11^

## Acknowledgments

D.L. Tharp (swine studies), S.M. Possidento and M.G. Turcotte (cell-based experiments), J. Li (snRNA-seq, RNA-seq, and protein-protein interaction studies), A.L. Bayer (snRNA-seq and T-cell analyses), A. Amin, P.K. Thorne and E. Wagoner (swine studies), F. Cividini (molecular analyses), C.L. Murray (mass spectrometry), X. Li (histology) and Y. Zhu (snRNA-seq): data collection and experiment analysis; R.V. Nair and V.B. Nguyen: bioinformatics; M.S. Kapiloff, C.A. Emter, D.L. Tharp, K.L. Dodge-Kafka, P. Alcaide and J.E. Van Eyk: project supervision, analysis and interpretation, and writing of manuscript. This work utilized bioinformatics services and computing resources provided by the Stanford Genetics Bioinformatics Service Center.

## Supplemental Material

Supplemental Methods

Major Resources Table

Figures S1-S11

Tables S1-S9 References 46-62

## Non-standard Abbreviations and Acronyms

AAV: adeno-associated virus
AAV9sc: self-complementary serotype 9 AAV vector
AKAP: A-kinase anchoring protein
CaMKII: Ca^2+^/calmodulin-dependent protein kinase II
CaMKAR: CaMKII activity reporter
CREB: cAMP Response Element Binding Protein
EDPVR: end-diastolic pressure volume relationship
eNOS: endothelial nitric oxide synthase
ERK: extracellular signal-regulated kinase
GFP: green fluorescent protein
HDAC4: histone deacetylase
4 HF: heart failure
HFpEF: heart failure with preserved ejection fraction
HFrEF: heart failure with reduced ejection fraction
HOMA-IR: homeostatic model assessment of insulin resistance
IκB: Inhibitor of NF-κB
IKKβ: IκB Kinase Subunit
β ITR: inverted terminal repeat
KBD: kinase binding domain
log_2_FC: log_2_ fold change
LV: left ventricle
NAb: neutralizing antibodies
NF-κB: nuclear factor κB
NE: norepinephrine
RNA-seq: RNA sequencing
PLN: phospholamban
RSK3: ribosomal S6 kinase type 3
snRNA-seq: single nucleus RNA sequencing
SRF: serum response factor
TNNT2: cardiac troponin T promoter
XBP1s: spliced X-box binding protein 1
WPRE: Woodchuck Hepatitis Virus posttranscriptional regulatory element
vg: viral genomes

